# netMUG: a novel network-guided multi-view clustering workflow for dissecting genetic and facial heterogeneity

**DOI:** 10.1101/2023.05.04.539350

**Authors:** Zuqi Li, Federico Melograna, Hanne Hoskens, Diane Duroux, Mary L. Marazita, Susan Walsh, Seth M. Weinberg, Mark D. Shriver, Bertram Müller-Myhsok, Peter Claes, Kristel Van Steen

**Author notes:** **Corresponding author:** Zuqi Li.

## Abstract

Multi-view data offer advantages over single-view data for characterizing individuals, which is crucial in precision medicine toward personalized prevention, diagnosis, or treatment follow-up. Here, we develop a network-guided multi-view clustering framework named netMUG to identify actionable subgroups of individuals. This pipeline first adopts sparse multiple canonical correlation analysis to select multi-view features possibly informed by extraneous data, which are then used to construct individual-specific networks (ISNs). Finally, the individual subtypes are automatically derived by hierarchical clustering on these network representations. We applied netMUG to a dataset containing genomic data and facial images to obtain BMI-informed multi-view strata and showed how it could be used for a refined obesity characterization. Benchmark analysis of netMUG on synthetic data with known strata of individuals indicated its superior performance compared with both baseline and benchmark methods for multi-view clustering. In addition, the real-data analysis revealed subgroups strongly linked to BMI and genetic and facial determinants of these classes. NetMUG provides a powerful strategy, exploiting individual-specific networks to identify meaningful and actionable strata. Moreover, the implementation is easy to generalize to accommodate heterogeneous data sources or highlight data structures.

**Author summary:** In recent years, we see the increasing possibility of collecting data from multiple modalities in various fields, requesting novel methods to exploit the consensus among different data types. As exemplified in systems biology or epistasis analyses, the interactions between features may contain more information than the features themselves, thereby necessitating the use of feature networks. Furthermore, in real-life scenarios, subjects, such as patients or individuals, may originate from diverse populations, which underscores the importance of subtyping or clustering these subjects to account for their heterogeneity. In this study, we present a novel pipeline for selecting the most relevant features from multiple data types, constructing a feature network for each subject, and obtaining a subgrouping of samples informed by a phenotype of interest. We validated our method on synthetic data and demonstrated its superiority over several state-of-the-art multi-view clustering approaches. Additionally, we applied our method to a real-life, large-scale dataset of genomic data and facial images, where it effectively identified a meaningful BMI subtyping that complemented existing BMI categories and offered new biological insights. Our proposed method has wide applicability to complex multi-view or multi-omics datasets for tasks such as disease subtyping or personalized medicine.

## Introduction

In machine learning, unsupervised clustering has been widely discussed and applied. It refers to a data-analysis problem where the true classification of individuals (or items) is unknown, and we derive clusters by exploiting between-individual similarity. Patient subgrouping plays an essential role in precision medicine, given that traditional medicine tends to offer a one-size-fits-all solution over the entire population and often overlooks heterogeneity [1,2]. While designing a drug customized for every patient may not be feasible, fine-scaled disease subtyping is tractable and can facilitate more personalized prevention, diagnosis, and treatment.

To better characterize a disease or phenotype, it is beneficial to turn to different sources, i.e. multi-view data, that are jointly more comprehensive and informative than single modalities [3]. For example, prior work has shown that multi-view clustering algorithms often outperform single-view ones [4–6]. However, many multi-view clustering methods have difficulty finding the consensus between modalities or exploiting relationships within and between views. Canonical correlation analysis (CCA) provides a solution to obtain optimal linear transformations of every data type to maximize their correlation [7]. Morover, sparse CCA (sCCA) introduces a sparsity parameter for each view which can enforce the canonical weights on most features to be zero [8]. sCCA both reduces the feature dimensionality and removes noisy features. Therefore, we can use this method to select the most meaningful features from datasets as input for the subsequent clustering. Sparse multiple canonical correlation network analysis (SmCCNet) can take an extraneous variable to detect modules of multi-view features with maximal canonical correlation between the data views informed by a phenotype of interest [9].

Current multi-view sample clustering methods integrate data views in different ways, e.g. concatenating all features, mapping views to a shared low-dimensional space, and merging between-sample relationship matrices from every view. The approach iCluster+ was designed to predict cancer subtypes from various data types by constructing a common latent space from all views [10]. The extension iClusterBayes adopts a Bayesian model in the latent space [11]. Other state-of-the-art methods focus on combining similarity matrices from every data view. For example, Spectrum is such a method with a self-tuning density-aware kernel and similarity network fusion (SNF) transforms data views to sample networks and fuses them nonlinearly [12,13]. However, these methods either reform the feature space or compute the between-sample interactions from features, which becomes deficient with data containing much information in the between-feature interactions.

Feature interactions derived from large sample collections are not individual-specific, albeit this has been shown to highlight interwiring modules for risk prediction and corresponding sample subtyping. Some analysis flows that illustrate this are built on the weighted gene co-expression network analysis (WGCNA) algorithm [14]. Starting from global networks, namely networks built on a collection of samples, each individual’s perturbation to the global network can be used to derive a so-called individual-specific network or ISN [15]. Comparing ISNs thus implies comparing interaction profiles between individuals. In reality, different features contributing to individual wirings may have different origins, paving the way toward system-level clustering of individuals in a precision medicine context.

In this work, we propose netMUG (**net**work-guided **MU**lti-view clusterin**G**), a novel multi-view clustering pipeline that clusters samples on the basis of these sample-specific feature interactions or wirings (Fig 1): In the presence of 2-view data, multi-view features are jointly selected via SmCCNet, based on canonical correlations and an additional extraneous variable. ISNs are constructed from the selected features, taking as a measure of edge strength the overall correlation between every pair of features and the extraneous data. The Euclidean distance metric representing dissimilarities between ISNs is fed into Ward’s hierarchical clustering [16], using the R library Dynamic Tree Cut to automatically derive the number of clusters [17]. After evaluating netMUG’s performance on synthetic data, we applied our workflow to a collection of participants of recent European ancestry with both genotypic information and 3D facial images available [18]. The aim was to dissect and interpret between-individual heterogeneity, informed by BMI as extraneous data. The application showed the potential of netMUG to refine existing classifications for obesity and to identify novel gene-BMI associations not yet discovered by genome-wide association studies (GWAS).

**Fig 1.**
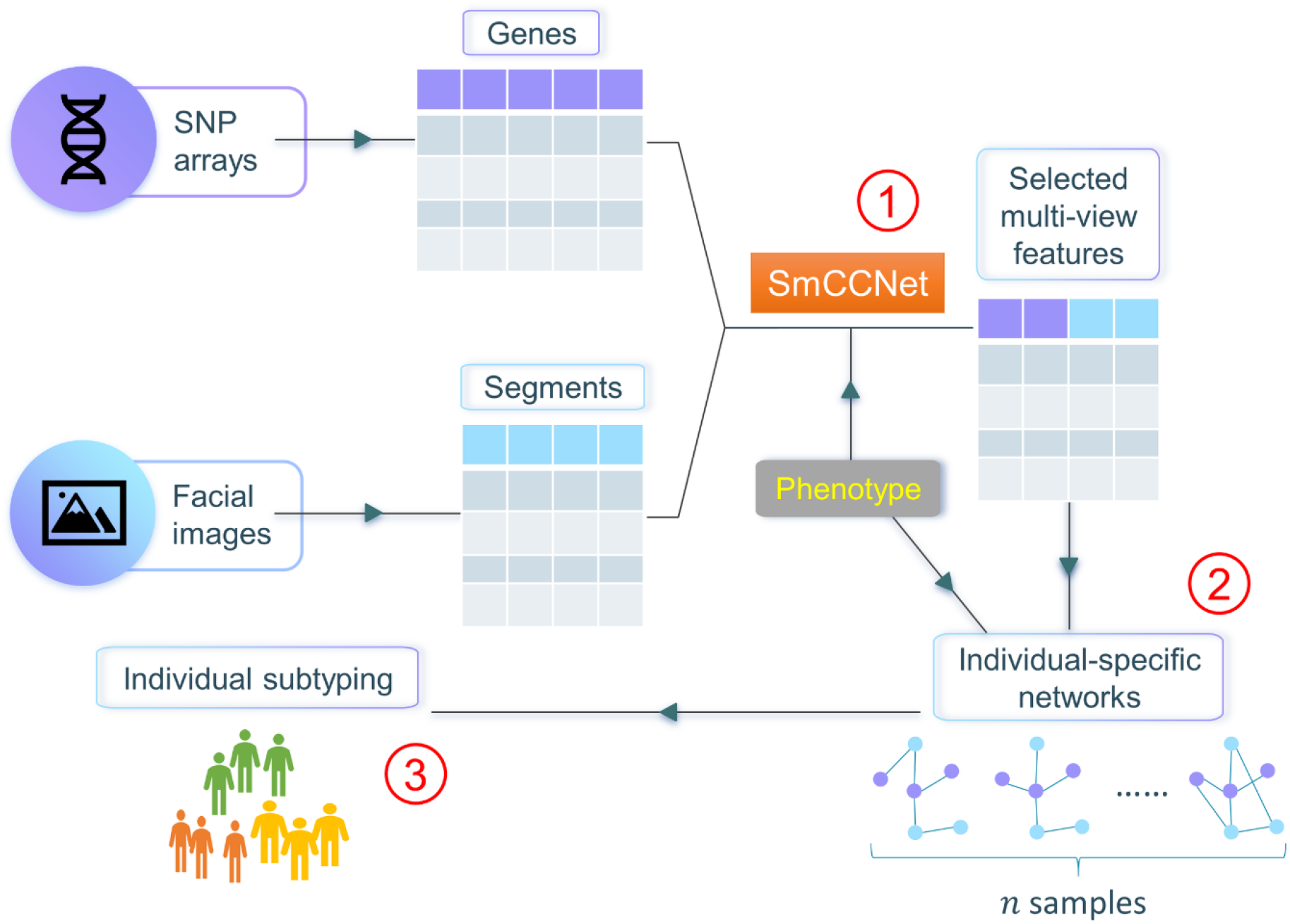
Workflow of netMUG (network-guided multi-view clustering). The pipeline consists of three parts: ① select phenotype-informed features from multiple data modalities via SmCCNet; ② build individual-specific networks based on the selected features and the phenotypic information; ③ subtype individuals via Ward’s hierarchical clustering.

## Materials and Methods

The code of netMUG is available on GitHub (https://github.com/ZuqiLi/netMUG.git). The complete pipeline, simulation, and analyses were done in R (version 4.2.1). The data representation of the case study was computed in Python (version 3.9.7).

For the remainder of this paper, we denote the two data views as *X* ∈ ℝ^*n*×*p*^ and *Y* ∈ ℝ^*n*×*q*^, respectively, the phenotypic variable as *Z* ∈ ℝ^*n*^, where *n* is the number of individuals, *p* the number of features in view 1, and *q* the number of features in view 2.

### SmCCNet

SmCCNet pipeline consists of a sparse multiple canonical correlation analysis (SmCCA) and a module detection method. CCA and its variants are a set of multivariate statistical methods based on the cross-covariance matrix. The basic CCA aims to find a pair of vectors *a* ∈ ℝ^*p*^ and *b* ∈ ℝ^*q*^ that maximizes the correlation between *Xa* and *Yb*:

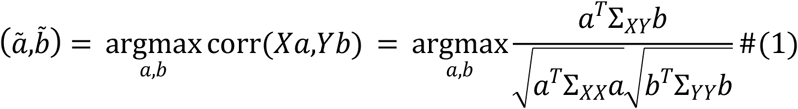

where ∑_*XY*_ denotes the cross-covariance matrix between *X* and *Y*, ∑_*XX*_ and ∑_*YY*_ are the covariance matrix of *X* and *Y*, respectively. If we standardize both *X* and *Y* so that every feature has a mean of 0 and a standard deviation of 1, Formula (1) can be reduced to:

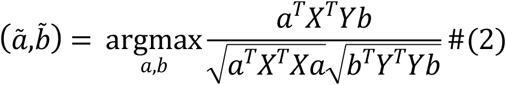

If we further constrain the covariance matrices to be diagonal, Formula (2) can be rewrite as:

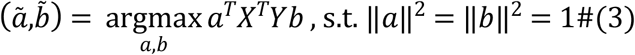

In case there are a lot of features with very little contribution to the canonical correlation, a sparse version of CCA (sCCA) is introduced, which applies the *ℓ*_1_ regularization to the canonical weights *a* and *b*. Hence the objective of sCCA is as follows:

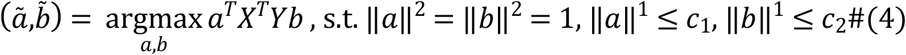

, where *c*_1_ and *c*_2_ are user-defined constants that regulate the sparsity, and 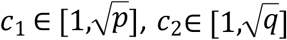. Their values can be chosen via cross-validation.

CCA and sCCA are originally designed for only two data views, however, additional information may be available and it may be helpful to take into account the correlations between the existing views and the extra one. In the special case where the new data type contains only a single feature, this can be considered as finding the optimal canonical pair that also corelates with the phenotype of interest. With the phenotypic variable denoted as *Z* ∈ ℝ^*n*^, the objective of SmCCA becomes:

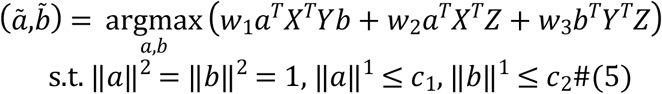

Coefficients *w* _1_, *w*_2_, *w*_3_ balance the three canonical correlations to account for the different correlation strength among the multiple data types.

The pair of canonical weight vectors 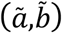 learnt from the SmCCA are sparse indicator of how much every feature in *X* and *Y* contributes to the overall correlation among the two data modalities and the phenotype. If we concatenate *ã* and 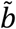 to form a new vector 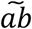, the outer product of 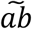 and itself gives us a similarity matrix whose elements measure the relatedness between every two features:

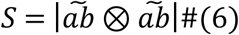

, where ⊗ is the outer product operator and | · | takes the absolute value of every element in a matrix.

To make the canonical weights robust, SmCCNet integrates a feature subsampling scheme which results in multiple pairs of 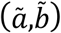 with different subsets of features from *X* and *Y*. The similarity matrices from each pair are then averaged and divided by the maximum.

Hierarchical clustering with complete linkage is performed on the distance matrix 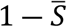, where 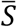 denotes the averaged and rescaled similarity matrix. The dendrogram is subsequently cut at the user-specified height and modules with feature(s) from a single view are discarded to derive multi-view modules.

### netMUG: workflow and algorithm

The input of netMUG is multi-view data with a phenotype (or more generally extraneous variables), both describing the same set of samples. To reduce dimensionality and extract the most informative features, netMUG first incorporates SmCCNet for feature selection, namely features in the final modules detected by SmCCNet. We have found that in practice, it is difficult to assess and quantify the balancing weights on each pairwise correlation (see in Formula (5)), so this property has been omitted in netMUG, i.e. *w*_1_ = *w*_2_ = *w*_3_ = 1.

We then use the selected features to construct ISNs for each individual. These networks are characterized by individual-specific edges as well as individual-specific nodes. In particular, we first construct a global network *G*^(*α*)^ = (*V,E* ^(*α*)^) across all samples, where *V* denotes the set of selected features from both views and *E*^(*α*)^ the set of edge weights whose element 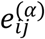 is the sum of pairwise Pearson correlations between feature *i, j* and the phenotype *Z* on all samples:

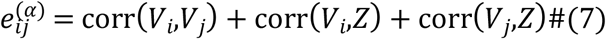

Second, we compute the leave-one-out network *G*^(*α*−*s*)^ = (*V,E*^(*α*−*s*)^) whose edges *E*^(*α*−*s*)^ are derived from all samples but sample *s* similarly to (3). Intuitively, the difference between the global and leave-one-out networks measures the perturbation on the network caused by sample *s*. Based on this, in our work, we define ISN by the absolute differential network:

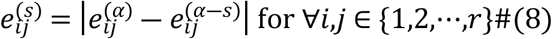

where 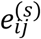 is the edge value between node *i* and *j* in the network specific to individual *s*. Hence, *G*^(*s*)^ = (*V,E*^(*s*)^), and *r* is the number of selected features. Formula (8) is a variant of the original ISN construction method in [15] because deriving an individual-specific association is not essential when the final aim is to identify “distances” between individuals. A higher 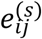 indicates that individual *s* deviates from its population more than others concerning the joint correlation between features and phenotype. In addition, an ISN with patterns very different from another means that their corresponding individuals may belong to different population subgroups.

The edge weights in an ISN can be seen as the realization in an *k*-dimensional Euclidean space with *k* = number of edges. Accordingly, clusters can be achieved by hierarchical clustering on Euclidean distance and Ward’s linkage. Because the Euclidean distance is not squared, we choose ‘ward.D2’ as method for the R function ‘hclust’ to adopt the correct Ward’s clustering criterion[19]. The clusters are obtained automatically by cutting the hierarchical tree at a height determined by the R package ‘dynamicTreeCut’ with deepSplit=1 to have fewer but larger clusters.

### Simulation study

We simulated 1000 samples with two views, each containing 2000 variables. Samples are randomly distributed in three balanced clusters. Out of the 2000 features, 200 are correlated between the two views with random covariance ranging from 0.5 to 0.8, i.e. [*X*_*corr*_ *Y*_*corr*_]∼*N* (0,*∑*) where 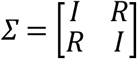 is an identity matrix and *R* ∈ ℝ^200×200^ is a diagonal matrix with the 200 covariances on the diagonal. There are another 200 features in each view, *X*_*clust*_ and *Y*_*clust*_, generated similarly but with covariance ranging from 0.8 to 1. These features are distinctive among the three sample clusters by multiplying 1,2,3 by their values for the samples in every cluster. The clustering also gave rise to a phenotype that follows a different normal distribution in every cluster, i.e. *Z* = [*Z*^(1)^ *Z*^(2)^ *Z*^(3)^]^*T*^ where *Z*^(1)^∼*N*(−1,1), *Z*^(2)^∼*N* (0,1), *Z*^(3)^∼*N*(1,1). To make the correlation patterns more complex and representative of real-life data, we linearly transformed [*X*_*corr*_ *X*_*clust*_] and [*Y*_*corr*_ *Y*_*clust*_]. The transformation coefficients were randomly sampled from the range [− 3,3]. Then 600 uncorrelated features per view all follow the standard normal distribution *N*(0,1), representing the noise.

We also investigated the execution time spent by each model. The bottleneck of netMUG is the SmCCNet step that computes a full SVD internally for each dataset, whose time complexity is 𝒪(min (*mn*^2^,*nm*^2^)) where *n* and *m* are the two dimensions of the data matrix. To make the method applicable to large-scale datasets, e.g. the whole genomic data composed of millions of SNPs, we may replace the full SVD with truncated SVD in the future for much faster computation and less memory usage.

### Case study

#### Data representation

Data pre-processing resulted in 265 277 SNPs and 7160 3D facial landmarks for all 4680 individuals with European ancestry (detailed steps are described in Supplementary). We subsequently reduced the dimensionality of both genomic and facial data via principal component analysis (PCA). Specifically, SNPs were first mapped to protein-coding genes if they fell within 2000 base pairs around a gene. Genes mapped with less than three SNPs were removed for insufficient information. Then, we performed PCA on the SNPs of every gene to obtain the principal components (PCs) explaining at least 90% of its variance. Meanwhile, we hierarchically segmented 3D facial images into five levels and 63 segments via the method proposed in [18]. Finally, the number of PCs representing every facial segment was determined via the simulation-based parallel analysis. As a result, we obtained 60 731 PCs for 9077 genes and 1163 PCs for 63 facial segments. In addition, more complex component summaries were also considered via diffusion kernel PCA [20], which nonlinearly exploits the graphic structure in the data. However, it is computationally intensive and requires extra hyperparameter tuning, and hence was discarded in this study.

#### Group-level interpretation

To estimate the association of BMI with our genomic data, we first conducted a GWAS via PLINK to calculate the P-value of the Wald test (detailed steps are described in Supplementary), which would tell if a SNP is significantly associated with BMI. Only SNPs with FDR-adjusted P-values < 0.05 were kept and then mapped to genes, following the same mapping strategy as in the ‘Data representation’ step. Meanwhile, we also downloaded the list of BMI-relevant genes from the DisGeNET database with a decent filter (>= 0.1) on the gene-disease association (GDA) score.

To further analyze the behavior of SmCCNet, we reran it with the same settings but without phenotypic information (the two sparsity parameters were re-determined via cross-validation). So far, four sets of genes have been obtained from the standard SmCCNet, GWAS, DisGeNET, and the uninformed SmCCNet. We investigated the overlap among them to see the agreement between SmCCNet and GWAS, the enrichment of DisGeNET genes in the gene set of SmCCNet (detailed steps are described in Supplementary), and the difference in SmCCNet made by the presence of phenotype.

#### Cluster-level interpretation

We first evaluated the association between the clustering derived by our method and BMI via a Kruskal-Wallis test. Our obesity subtypes were also compared with the classic BMI categories, i.e. underweight (BMI < 18.5), normal (18.5 <= BMI < 25), overweight (25 <= BMI < 30), and obese (BMI >= 30) [21].

Further, we characterized every subgroup by facial and genetic information. By averaging all facial images of each subgroup, we represented it with a mean face shape. As for genetics, ISNs of every cluster were averaged, and a subnetwork was taken from the mean ISN based on the top 1% edge weights to only focus on the vital signals. Subsequently, we computed the largest connected component of every subnetwork and extracted all genes in this component. The overlap among clusters was analyzed, and we gave special attention to the cluster-specific genes to characterize each cluster. In particular, an enrichment analysis was done via the analysis tool of the Reactome website [22] to test which biological pathways are significantly over-represented in the gene list specific to every cluster.

#### ISN-level interpretation

Because our pipeline represents every individual by a network, namely ISN, a fully-connected weighted graph, we need a lower-dimensional representation of the ISNs to examine their behaviour better and visualize them. Therefore, the graph filtration curve method was applied, which computed a function or curve from each ISN, whose values were derived via graph evolution [23]. More specifically, we set a series of thresholds on the edge weights and, at each threshold, constructed a subnetwork by taking only edges larger than that threshold. In such a manner, we got a series of subnetworks from smallest to largest for every ISN, and the largest subnetwork is the ISN itself. Then some graph property was calculated based on each subnetwork, and therefore, an ISN was converted to a function of graph property against edge threshold. Here in our project, we chose the mean node degree of the largest connected component (LCC) as the graph property because the subgraphs may not be fully connected anymore. The average degree is a simple yet powerful tool to measure graph density.

## Results

netMUG was validated on a synthetic dataset and compared with both baseline and benchmark methods for multi-view clustering. We then applied it to real-life large-scale multi-view cohort data and characterized the resultant clusters by their representative faces and enriched pathways.

### Validating netMUG with synthetic data

We simulated a scenario where a multi-view dataset contained complex feature patterns and many noisy features, representing real-life high-dimensional data, e.g. genomics and images. Two criteria were adopted to assess clustering performance, P-value from the Kruskal-Wallis test and adjusted Rand index (ARI). Kruskal-Wallis test was used as a non-parametric version of one-way ANOVA to test whether the distribution of an extraneous phenotype is similar across clusters [24]. Therefore, a low P-value (< 0.05) indicates a significant association between the clustering and the phenotype. ARI was computed to show the similarity between the derived clustering and the ground-truth [25]: the higher, the better; an ARI of 1 means perfect fitting.

Various baseline and benchmark methods were considered in addition to netMUG, and their performances were compared. Baseline models included *k*-means on every single view, and both views concatenated, with or without principal component analysis (PCA) as data dimensionality reduction strategy. We chose four benchmark models, iCluster+ [10], iClusterBayes [11], Spectrum [12], and SNF [13], and their codes are available in R [26]. To show the effectiveness of SmCCNet as a feature selector we also applied Spectrum and SNF on the features selected by SmCCNet, i.e. model ‘SmCCNet-Spectrum’ and ‘SmCCNet-SNF’. Finally, as an alternative to our proposed framework, we replaced the hierarchical clustering with spectral clustering as the last step of netMUG, leading to the model ‘SmCCNet-ISN-SC’.

As shown in Fig 2 (exact values are listed in Supp. Table 1), all baseline models performed poorly in terms of ARI. The P-values for models with PCA on single *X* and both views were lower than 0.05, but did not convey a big difference on the plot. Among the benchmarks, SNF outperformed iCluster+, iClusterBayes, and Spectrum. With features selected by SmCCNet, the performances of both Spectrum and SNF were substantially improved, implying the necessity of feature selection on high-dimensional and noisy data. Integrating SmCCNet, ISN, and Dynamic Tree Cut, netMUG achieved the best P-value and ARI results. It retrieved the ground-truth clustering for the complex multi-view data we simulated.

**Fig 2.**
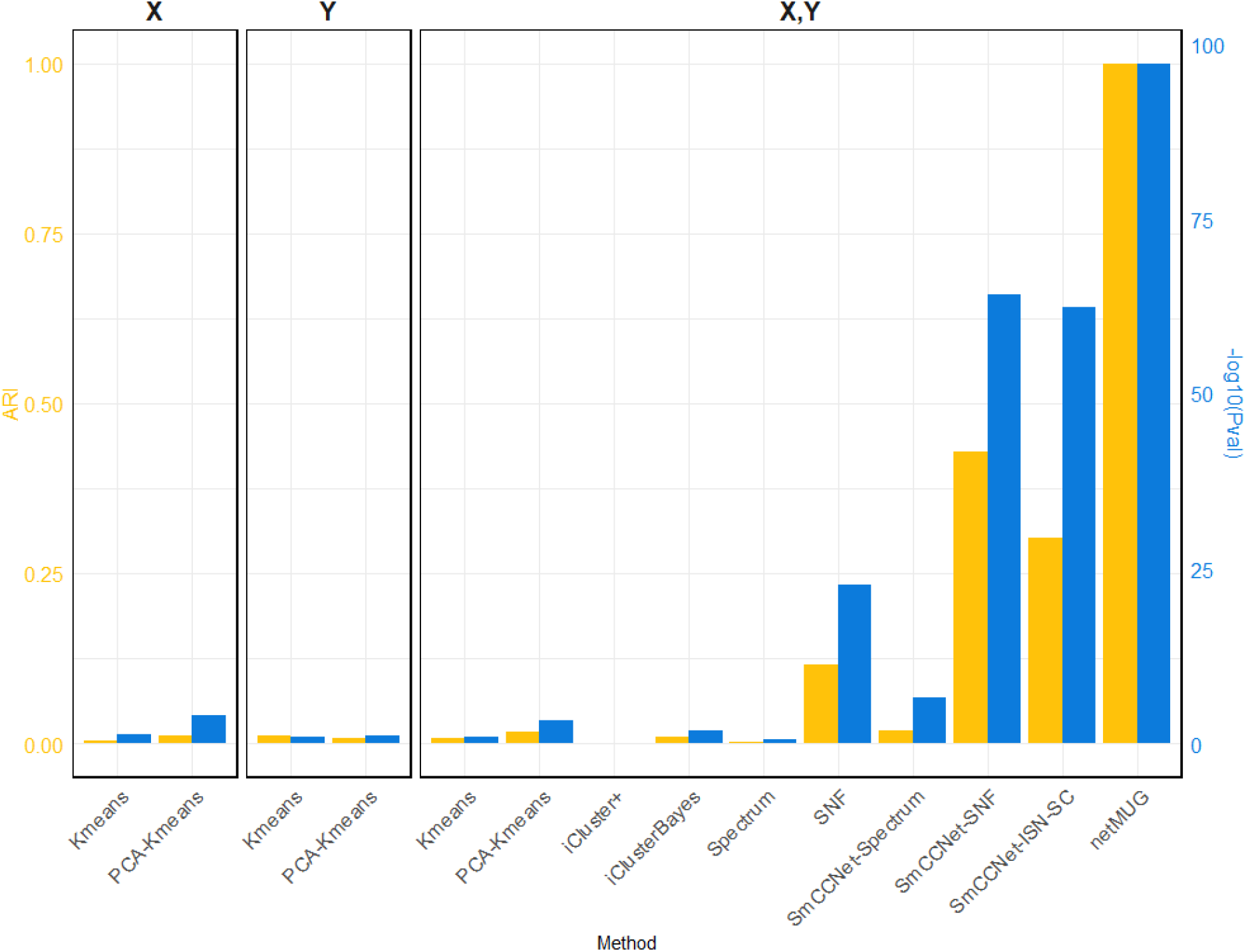
Performance of all methods on simulated data. The three barplot panels illustrate the ARI (on the left axis in yellow) and − log_10_ *Pval* (on the right axis in blue) for the four single-view (X or Y) and 10 multi-view (both X and Y) models. Four out of six baseline models (‘Kmeans’ and ‘PCA-Kmeans’) use a single view (X or Y). Two-view ‘Kmeans’ and ‘PCA-Kmeans’ models are based on the concatenation of X and Y. ‘SmCCNet-Spectrum’ and ‘SmCCNet-SNF’ are benchmark models with features selected by SmCCNet. ‘SmCCNet-ISN-SC’ only differs from the proposed framework, netMUG, for the clustering method.

The runtime of each method (Supp. Table 1) shows that all baseline models spent less than half minute and PCA brought an acceleration in speed for fewer features. iCluster+ and iClusterBayes cost much longer time than Spectrum and SNF without an improvement in performance. The SmCCNet feature selection step took 33 min on its own, which became the bottleneck of all the SmCCNet-based models (‘SmCCNet-Spectrum’, ‘SmCCNet-SNF’, ‘SmCCNet-ISN-SC’ and netMUG). This is mostly due to the subsampling scheme and the full SVD computation within SmCCNet.

### Case study

To exemplify netMUG, we used a multi-view dataset of 4680 individuals of recent European ancestry recruited from three independent studies in the US: 3D Facial Norms cohort (PITT), Pennsylvania State University (PSU) and Indiana University-Purdue University Indianapolis (IUPUI) [18]. For each individual, facial images and genomic data were collected along with extra information, including age, sex, height, and weight. BMI was computed as: BMI = weight(*kg*) height(*m*)^2^.

We interpreted netMUG analysis results at two levels: all-samples level, hereafter referred to as group-level, and at the level of clustered individuals. Group-level interpretation refers to describing the multi-view features selected by SmCCNet. In addition, we assessed the overlap between SmCCNet-selected genes and genes found by a genome-wide association study (GWAS) or DisGeNET database. Cluster-level interpretations were made by evaluating the association between the final clustering and BMI and the characterization of every cluster in terms of facial or genetic characteristics. Finally, we applied graph filtration curves to represent and visualize ISNs in 2D space [23].

Two flavors of netMUG were implemented: one with SmCCNet informed by BMI as extraneous variable, and one without such information.

#### Group-level interpretation

##### Informed by BMI

We chose the two sparsity parameters for SmCCNet, namely *c*_1_ and *c*_2_ in Formula (5), via 5-fold cross-validation. The subsampling proportions were determined to be 50% and 90% for genomic and facial data because of their substantial dimensional differences. We then computed the average canonical weights over 500 subsampling runs to derive the similarity matrix 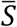. Three modules with 316 PCs (see Methods “Data representation”) were found by cutting the hierarchical tree at a height very close to 1 (0.999) and discarding modules with a single feature or features from a single view. The cutting threshold was determined following Shi WJ et al. [9] The selected features from the retained modules comprised 278 genes and 26 facial segments. All the subsequent analyses were performed on these features unless mentioned otherwise. One of the most well-known obesity-related genes, *FTO* [27], was in the gene list. Another essential gene for obesity, *MC4R* [28] was not selected because it was filtered out for having less than three SNPs mapped. A recent epistasis analysis found two pairs of SNPs whose interactions were associated with BMI (*FTO* – *MC4R* and *RHBDD1* – *MAPK1*) [29]. SmCCNet did not detect *RHBDD1* or *MAPK1*, possibly caused by CCA’s focus on inter-modal rather than intra-modal interactions.

GWAS detected 155 SNPs significantly associated with BMI mapped to 95 protein-coding genes (P-value < 0.05 adjusted by false discovery rate, or FDR). Out of these 95 genes, 16 (16.8%) were in common with the 278 genes selected by SmCCNet (Fig 3, set A and D), confirming the agreement between SmCCNet and GWAS but also explaining their difference in the involvement of facial information.

**Fig 3.**
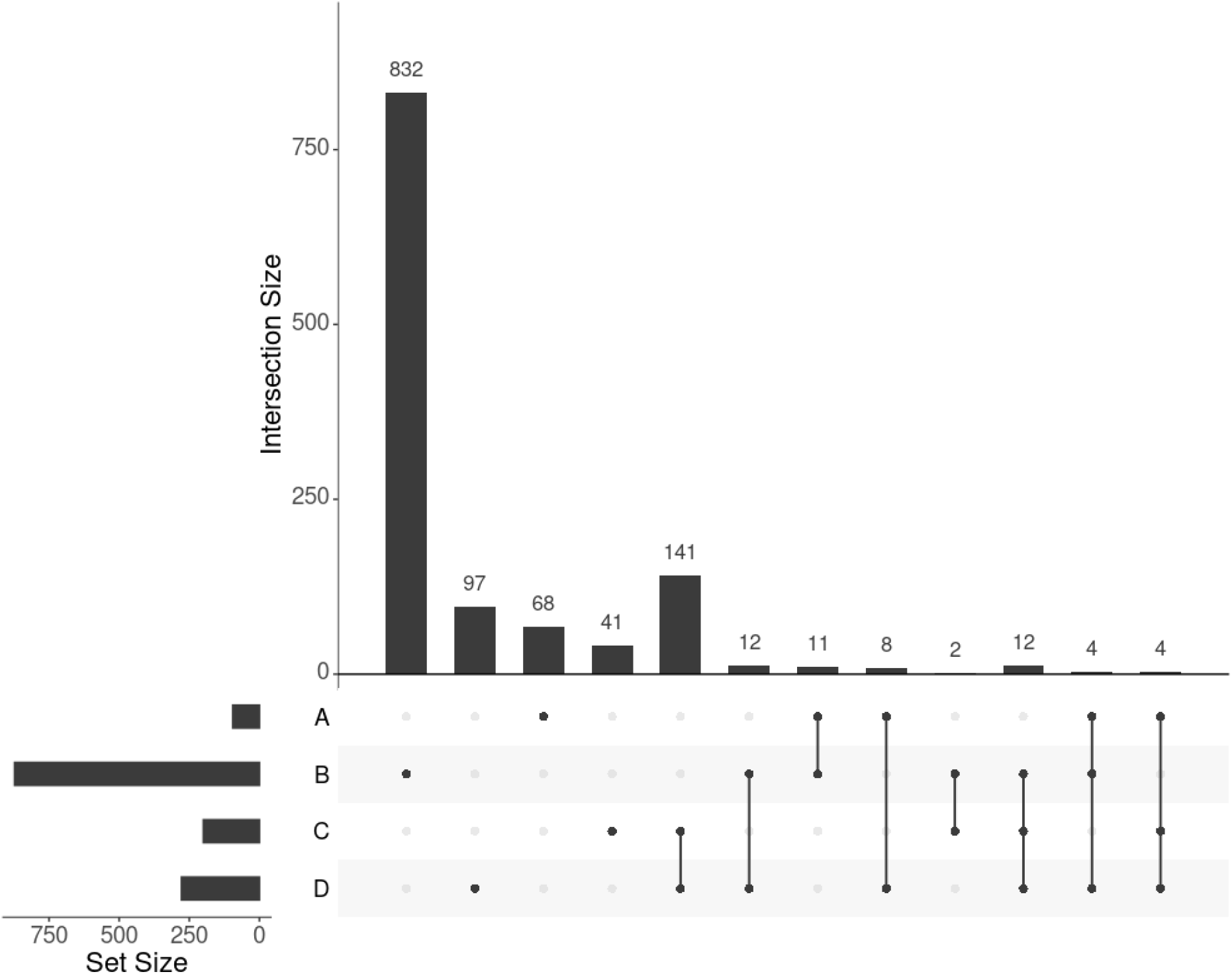
UpSet plot showing the intersections of four gene sets. A, B, C, and D represent the gene list found by GWAS, DisGeNET, uninformed SmCCNet, and standard SmCCNet, respectively. Uninformed SmCCNet maximizes the canonical correlation without extraneous information, whereas standard SmCCNet takes BMI into account to supervise CCA. Genes of the four sets are listed in Supplementary.

We found 1014 genes from DisGeNET associated with BMI with gene-disease association (GDA) score >= 0.1 [30], out of which 873 are protein-coding genes. Of the 873 genes, 28 (3.2%) also appear in the 278 genes selected by SmCCNet (Fig 3, set B and D). The hypergeometric test shows that the DisGeNET gene set is significantly enriched in the gene set from SmCCNet (P-value = 6.0 × 10^−5^).

##### Uninformed by BMI

A total of 329 features were selected by the uninformed SmCCNet, resulting in 200 genes and 50 facial segments. Of the 200 genes, 157 (78.5%) were shared between the standard and the uninformed SmCCNet (Fig 3, set C and D). However, the uninformed SmCCNet found fewer GWAS genes and DisGeNET genes than informed SmCCNet (Fig 3, set A, B, and C), highlighting the merits of supervised analysis.

We also looked at the top 1% of connections between genes and facial segments in the features selected by informed SmCCNet (Fig 4). It was clearly shown that the full face has high relatedness with all genes, indicating that those genes are primarily associated with facial morphology globally. On a local level, the selected genes are most strongly connected with the eyes (with the temporal area) and chin, in line with the fact that they are known facial signals for obesity. *AATK* and *CD226* are the genes with the most top connections. *AATK* plays a role in neuronal differentiation and has been known to be highly associated with BMI [31]. CD226 encodes a glycoprotein expressed on the surface of blood cells and is related to colorectal cancer [32], which could be a potential biomarker of obesity. Furthermore, *APOBEC3A, DNAJC5B*, and *NGFR* all affect body height and BMI [33].

**Fig 4.**
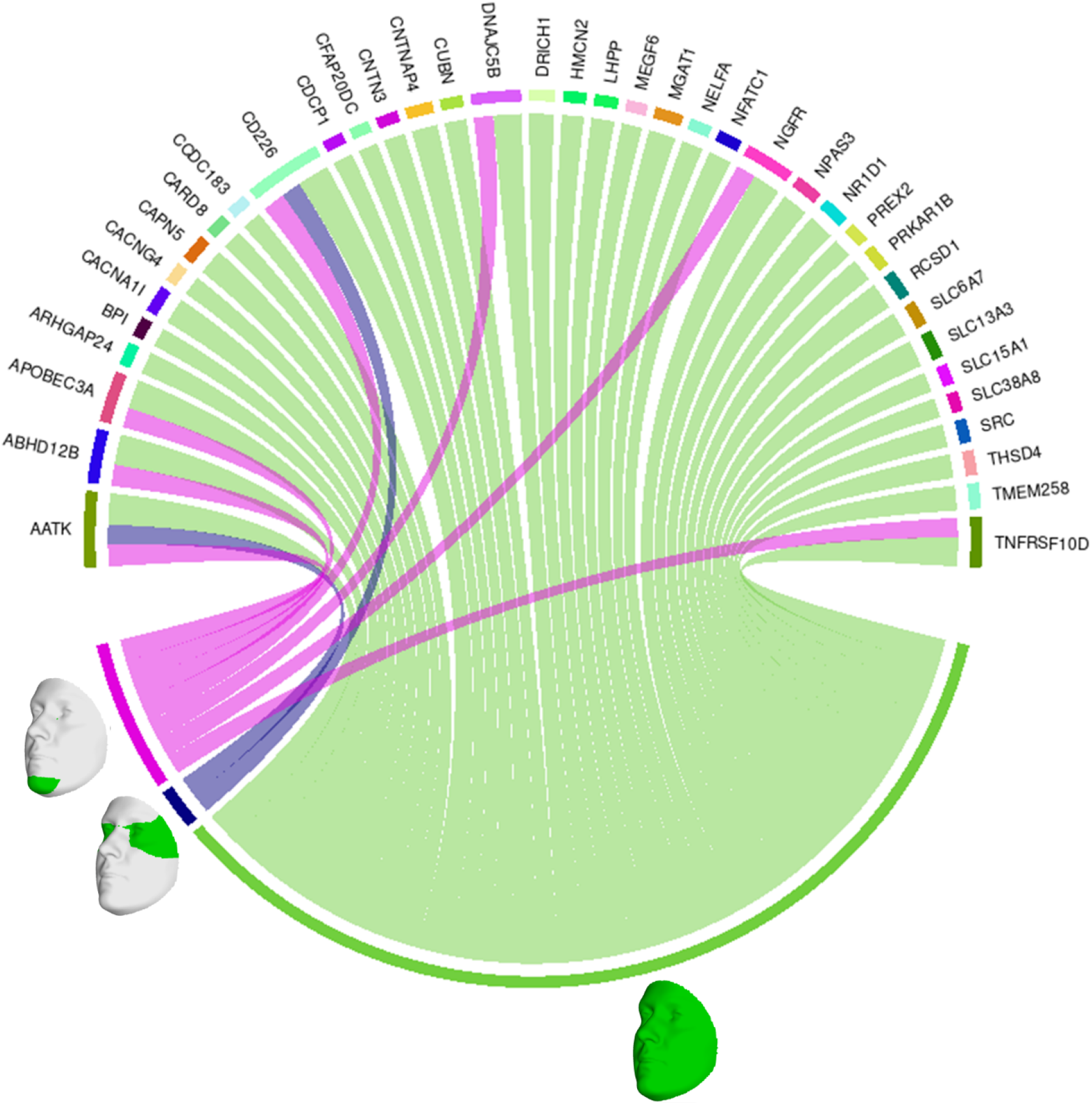
Genes and facial segments with the top 1% connections. Connections refer to the similarities between the canonical weights of genetic and facial PCs, as described in Methods. Only inter-modality relationships are considered. A thicker connection in the circus plot means a higher relatedness.

#### Cluster-level interpretation

netMUG automatically detected five clusters with significantly different BMI distributions given the Kruskal-Wallis test (P-value = 7.4 × 10^−150^) (Fig 5). Clusters 4 and 5 are the two outstanding groups, and both fall into the classic category of ‘Obese.’ [21] Nevertheless, as clearly shown in Fig 5, our clustering provides higher granularity on the obese group, which is desired because it can lead to more precise identification and better characterization of obese people. Cluster 1 contains roughly half people in the dataset, and it substantially follows the distribution of the whole population with most normal and normal-to-overweight individuals. Clusters 2 and 3 have similar bimodal distributions with different deviations. The two peaks of cluster 3 are farther away from the center than cluster 2. Jointly looking at clusters 1, 2, and 3, they classify individuals by how far they are from the ‘population normal’ while treating underweight and overweight indifferently. This behavior suggests that underweight and overweight people deviate from the normal condition similarly in terms of multi-view interactions, which may be related to, e.g. the double burden of malnutrition. It has been shown that some crucial vitamins and minerals can affect both underweight and overweight individuals.

**Fig 5.**
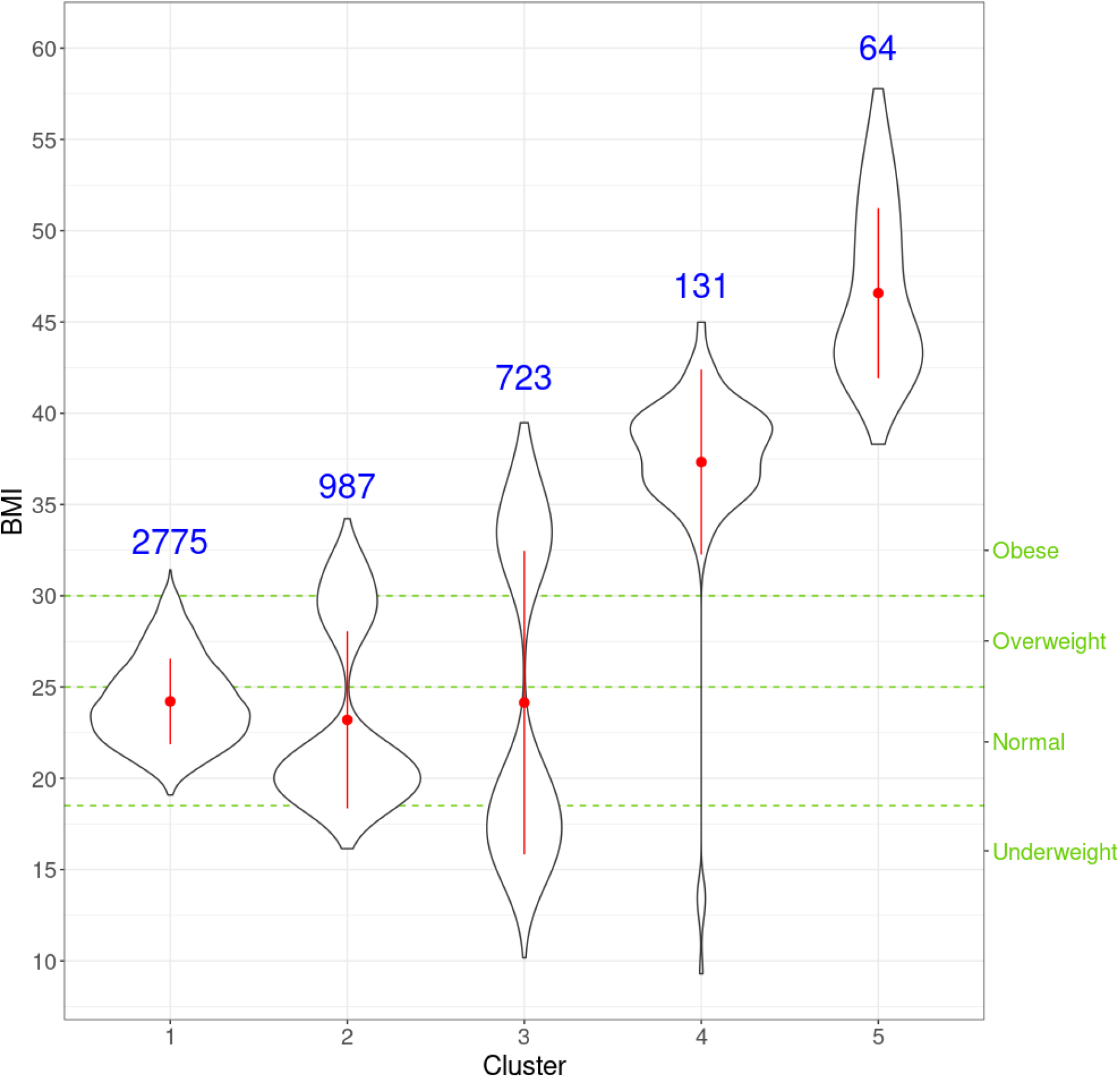
Violin plots for the distribution of BMI for individuals in every cluster. For each cluster, the red dot and vertical line indicate the mean and standard deviation, respectively, and the number in blue is the size of every cluster. The three green horizontal dashed lines represent the cut-offs of the four classic BMI categories shown on the right Y-axis.

Next, we look at genetic and facial characteristics of the identified clusters. The average facial shapes per cluster are depicted in Fig 6 and largely follow the profile of mean BMI across clusters (Fig 5). Again, clusters 4 and 5 stand out compared to clusters 1-3. Superposition of the average faces shows pronounced areas at the forehead or in the chin for cluster 4 individuals, whereas cheek and eye areas most responsible for cluster 5 differences from the rest of the samples. Cluster 1-3 faces have pronounced features around nose and mouth.

**Fig 6.**
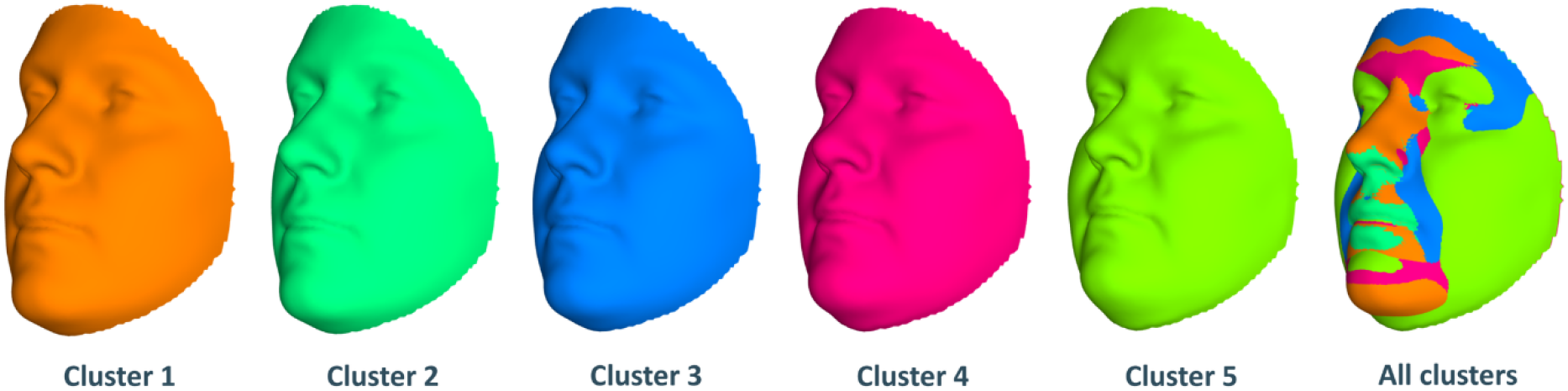
Mean facial shapes of every cluster and the superposition of them. The first five faces are the average of all individual faces in every cluster while the last face was obtained by plotting the five mean faces on top of each other (colors of this superposition face correspond to the five mean faces).

For individuals within every cluster, we averaged their corresponding ISNs and binarized the mean ISNs with top 1% edge weights being 1 and the rest 0. Subsequently, we computed the largest connected component (LCC) from every binary network, resulting in LCCs of 86, 129, 144, 112, and 136 nodes (68, 114, 119, 94, and 118 genes) for the five clusters, respectively (Fig 7). The genes *MIGA1, CACNA1B*, and *SLC38A8* were common to all clusters. *MIGA1* regulates mitochondrial fusion and shows a high relationship with BMI according to GWAS Catalog (scoring 14.0) [31]. *CACNA1B* encodes a voltage-gated calcium channel subunit and GWAS Catalog records a strong association (scoring 12.2) between *CACNA1B* and acute myeloid leukemia [34], for which BMI is a known risk factor. *SLC38A8* has a high GWAS Catalog score (14.1) with adiponectin measurement [35], which directly affects insulin sensitivity and obesity, and a strong association with eye diseases, e.g. foveal hypoplasia 2 and anterior segment dysgenesis. The link from gene *SLC38A8* to both obesity and facial features may imply a novel relationship between obesity and facial morphology.

**Fig 7.**
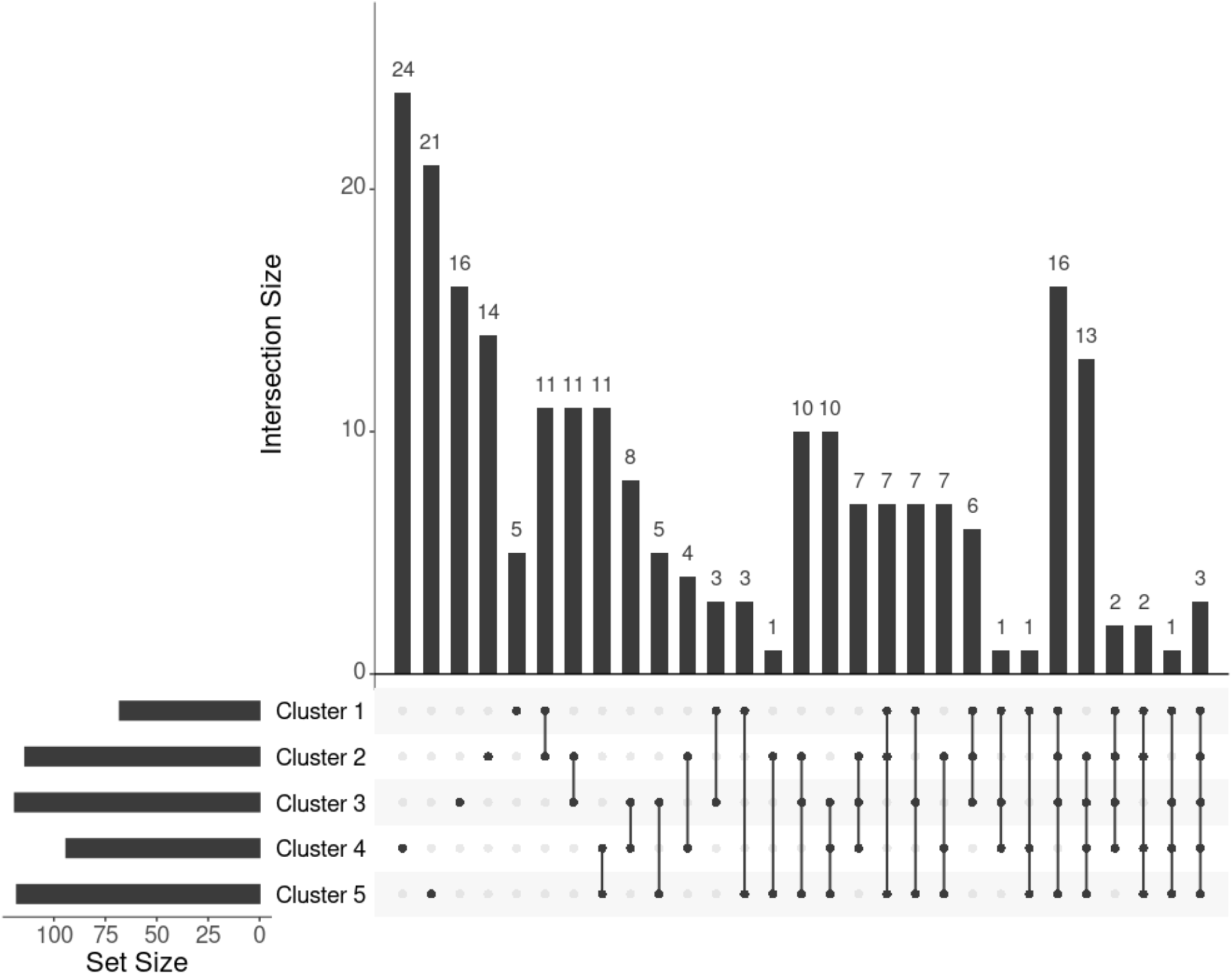
UpSet plot showing the intersections of genes extracted from the mean ISN of every cluster. To further investigate the genes exclusively extracted from each subgroup, i.e. subtype-specific genes, cluster 4 has the most subtype-specific genes (24) with also the highest proportion (25%) to all the genes in its LCC, followed by cluster 5 with 21 unique genes (17.8%). This observation is in line with the distinctive BMI distributions for these clusters. Meanwhile, there are only five genes specific to cluster 1 (7.4% of 68 genes in the LCC of cluster 1), suggesting that cluster 1 represents the ‘population normal.’

One or more Reactome pathways were significantly enriched in genes that were specific to a cluster, except for cluster 5 (Table 1). The reason may be that genes enriching cluster 5 were also obtained in other clusters, so they were not considered specific to cluster 5. Another possibility is that those 21 genes exclusively in cluster 5 are too functionally diverse as obesity is involved in many different biological pathways.

**Table 1.** Enriched Reactome pathways in every cluster. Pathways for cluster 5 are all in italic because no P-value was lower than 0.05 after multiple testing correction (FDR). For cluster 5, the four pathways with raw P-values ≤ 0.01 are shown.

|  |  |
| --- | --- |
| **Cluster 1** | Tryptophan catabolism |
| **Cluster 2** | TP53 regulates transcription of death receptors and ligands<br>Defective GALNT12 causes CRCS1<br>Defective GALNT3 causes HFTC<br>NPAS4 regulates expression of target genes<br>Transcriptional regulation by NPAS4 |
| **Cluster 3** | Regulation of TP53 expression |
|  | STAT3 nuclear events downstream of ALK signaling<br>Signaling by ALK<br>Regulation of TP53 expression and degradation |
| **Cluster 4** | Regulation of gene expression in early pancreatic precursor cells<br>Regulation of gene expression in late stage (branching morphogenesis) pancreatic bud precursor cells<br>Rap1 signalling<br>Defective B3GALTL causes PpS<br>O-glycosylation of TSR domain-containing proteins |
| **Cluster 5** | *Glycogen storage disease type Ib (SLC37A4)*<br>*Interleukin-33 signaling*<br>*RHOB GTPase cycle*<br>*RHOC GTPase cycle* |

#### ISN-level interpretation

With the mean node degree of the largest connected component being the function to describe a graph, the filtration curves of ISNs exhibited notable variation in their evolution trajectory, which implies the ability of ISN to exploit the between-individual heterogeneity (Supp. Fig 2). If we group the filtration curves by the netMUG-derived clustering, their values are significantly different at most edge thresholds (Fig 8). Clusters 4 and 5 have higher mean degree values overall than the rest along the graph evolution, indicating that their ISNs are depicted by more densely connected components. This result is in line with the association between BMI and every cluster (Fig 5) in the sense that people with severe obesity (clusters 4 and 5) will show more prominent characteristics than normal or slightly overweight people (clusters 1, 2, and 3). The ISN-level results both mean that average degree is a good descriptor for ISNs and confirm that the clusters differ in terms of intrinsic graph properties, e.g. average degree.

**Fig 8.**
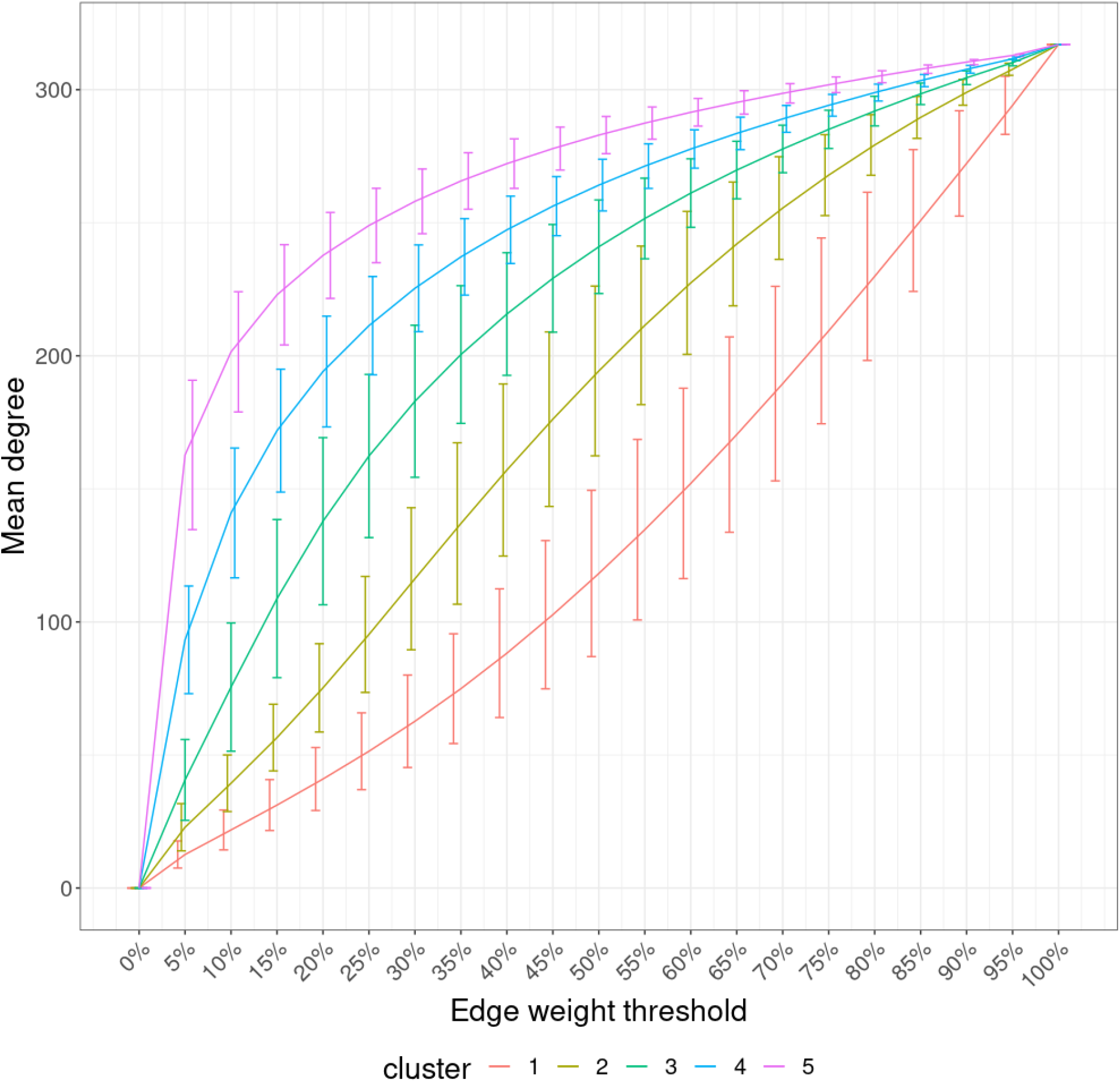
Filtration curves of ISNs grouped by the derived clustering. Every line linking the bottom-left and upper-right corners shows the mean filtration curve of each cluster. The vertical lines are the standard deviation of the function values at each threshold within each cluster.

## Discussion

In this study, we introduced netMUG, a multi-view clustering strategy that links population-based to individual-specific networks, possibly informed by extraneous variable. Synthetic data analysis showed promising outperformance of netMUG compared to baseline and benchmark multi-view clustering methods. An application to real-life cohort data, and exploiting extraneous information via BMI, revealed a refined classification of obese individuals and increased understanding about genetic and facial segments linked to BMI-induced subgroups.

Because our workflow is highly modular, various extensions or adaptions can be implemented, at the discretion of the user. Here, we presented a basic version. Modifications can be made at several levels. At the level of the data input, a single data view can be considered, a single or multiple extraneous variables can be informative, missing values can be inferred by imputation. Single-view netMUG replaces its basic SmCCNet implementation with one targeting a single dataset only. With multiple extraneous variables (for instance a symptom group of variables), the weight optimization (in Formula (5)) will be affected via prior knowledge or cross validation. When adopting imputation strategies, it is important to account for within- and between-data relationships. We refer to [36], who reviewed integrative imputation strategies for multi-omics datasets. Some of the proposed strategies may also apply to highly heterogeneous omics/non-omics mixed data. These include deep learning inspired approaches.

Also, adaptions are possible at the level of computing relationships between sets of variables. For instance, structural equation models (SEMs) can be used to conduct CCA in the presence of missing data [37]. It needs more work to see how sparsity and supervision are best introduced in SEMs as in sparse CCA, which was key to SmCCNet [9]. For a review of sparse CCA extension to genomic data, we refer to [38]. Then, ideas on sparse supervised CCA and sparse multiple CCA (more than two data views) pave the way for extended applications of netMUG, for instance via the R function MultiCCA [8].

Once a filtered heterogeneous network is obtained, its nodes can serve as template to construct individual-specific edges. It is up to the user to define the most appropriate measure of association. Our analyses on real-life datasets showed a benefit to include the extraneous data in this measure (here in this work: BMI). Clearly, the appropriateness of a measure of association between features will be context-dependent.

At the level of sample clustering, multiple graph clustering algorithm can be used. The diversity in algorithm is in part due to the many ways distance or similarity between graphs (here: ISNs) can be defined. Examples include edge difference distance or graph diffusion distance. These options and many more are discussed in [39]. The basic version of netMUG represents ISNs as points in a high-dimensional Euclidean space (axes refer to edges) and implements Ward’s minimum variance hierarchical clustering method. The initial cluster distance is taken to be the squared Euclidean distance between points.

Several measures can be taken to further increase the robustness of netMUG. Our choice is to adopt the feature subsampling in SmCCNet, i.e. SmCCA is run multiple times with each time on a random subset of features. It relates to increasing the robustness of feature selection. Robustness may also be increased at the clustering level by fusing a variety of clustering algorithms or settings within a multiple clustering protocol [40]. Deriving the optimal number of clusters via Dynamic Tree Cut settings or computationally more intensive multiscale bootstrap resampling may be included in such a protocol. Alternatively, the hierarchical clustering process and significance assessment are intertwined as in netANOVA [41], with promising performance for ISNs.

We illustrated netMUG on cohort data, with BMI as extraneous information. Alternative applications of netMUG include disease subtyping and studies that explore the impact of confounders on clustering results. The latter studies can be carried out by comparing supervised (i.e. with the confounders or extraneous information) and unsupervised netMUG (i.e. without *Z* in Formula (5)).

As conclusion, our proposed netMUG method exploits population-based and individual-specific network analyses to construct and interpret multi-view clusters. Clusters can or cannot be supervised by using extraneous data. The modular build-up of the workflow easily allows customization at several steps in the workflow, for instance going beyond two data views.

## Data availability statement

For the 3D Facial Norms dataset, genotypic markers are available to the research community through the dbGaP controlled-access repository (http://www.ncbi.nlm.nih.gov/gap) at accession #phs000929.v1.p1. The raw source data for the phenotypes - the 3D facial surface models in .obj format - are available through the FaceBase Consortium (https://www.facebase.org) at accession #FB00000491.01. Access to these 3D facial surface models requires proper institutional ethics approval and approval from the FaceBase data access committee. The PSU and IUPUI datasets were not collected with broad data sharing consent. The code of netMUG and simulated data are available on GitHub (https://github.com/ZuqiLi/netMUG.git).

## Author contributions

**Zuqi Li**: Conceptualization, Formal Analysis, Methodology, Validation, Writing – original draft. **Federico Melograna**: Methodology, Writing – review & editing. **Hanne Hoskens**: Methodology, Writing – review & editing. **Diane Duroux**: Methodology, Writing – review & editing. **Mary L. Marazita**: Resources, Writing – review & editing. **Susan Walsh**: Resources, Writing – review & editing. **Seth M. Weinberg**: Resources, Writing – review & editing. **Mark D. Shriver**: Resources, Writing – review & editing. **Bertram Müller-Myhsok**: Supervision, Writing – review & editing. **Peter Claes**: Resources, Supervision, Writing – review & editing. **Kristel Van Steen**: Conceptualization, Supervision, Writing – original draft, Writing – review & editing.

## Acknowledgements

Many thanks to the former PhD student Karlijne Indencleef at the department of Electrical Engineering, ESAT/PSI, KU Leuven, who greatly helped me understand how the US cohort data were pre-processed and how the imaging analysis was done.

## References

1. Spycher BD, Silverman M, Brooke AM, Minder CE, Kuehni CE. Distinguishing phenotypes of childhood wheeze and cough using latent class analysis. European Respiratory Journal. 2008;31: 974–981. doi:10.1183/09031936.00153507

2. Saria S, Goldenberg A. Subtyping: What It is and Its Role in Precision Medicine. IEEE Intell Syst. 2015;30: 70–75. doi:10.1109/MIS.2015.60

3. Gligorijević V, Malod-Dognin N, Pržulj N. Integrative methods for analyzing big data in precision medicine. Proteomics. 2016;16: 741–758. doi:10.1002/pmic.201500396

4. Chauvel C, Novoloaca A, Veyre P, Reynier F, Becker J. Evaluation of integrative clustering methods for the analysis of multi-omics data. Briefings in Bioinformatics. 2020;21: 541–552. doi:10.1093/bib/bbz015

5. Abavisani M, Patel VM. Deep Multimodal Subspace Clustering Networks. IEEE J Sel Top Signal Process. 2018;12: 1601–1614. doi:10.1109/JSTSP.2018.2875385

6. Wen Y, Song X, Yan B, Yang X, Wu L, Leng D, et al. Multi-dimensional data integration algorithm based on random walk with restart. BMC Bioinformatics. 2021;22: 97. doi:10.1186/s12859-021-04029-3

7. Hotelling H. RELATIONS BETWEEN TWO SETS OF VARIATES. Biometrika. 1936;28: 321–377. doi:10.1093/biomet/28.3-4.321

8. Witten DM, Tibshirani R, Hastie T. A penalized matrix decomposition, with applications to sparse principal components and canonical correlation analysis. Biostatistics. 2009;10: 515–534. doi:10.1093/biostatistics/kxp008

9. Shi WJ, Zhuang Y, Russell PH, Hobbs BD, Parker MM, Castaldi PJ, et al. Unsupervised discovery of phenotype-specific multi-omics networks. Bioinformatics. 2019;35: 4336–4343. doi:10.1093/bioinformatics/btz226

10. Mo Q, Wang S, Seshan VE, Olshen AB, Schultz N, Sander C, et al. Pattern discovery and cancer gene identification in integrated cancer genomic data. Proc Natl Acad Sci USA. 2013;110: 4245–4250. doi:10.1073/pnas.1208949110

11. Mo Q, Shen R, Guo C, Vannucci M, Chan KS, Hilsenbeck SG. A fully Bayesian latent variable model for integrative clustering analysis of multi-type omics data. Biostatistics. 2018;19: 71–86. doi:10.1093/biostatistics/kxx017

12. John CR, Watson D, Barnes MR, Pitzalis C, Lewis MJ. Spectrum: fast density-aware spectral clustering for single and multi-omic data. Cowen L, editor. Bioinformatics. 2019; btz704. doi:10.1093/bioinformatics/btz704

13. Wang B, Mezlini AM, Demir F, Fiume M, Tu Z, Brudno M, et al. Similarity network fusion for aggregating data types on a genomic scale. Nat Methods. 2014;11: 333–337. doi:10.1038/nmeth.2810

14. Langfelder P, Horvath S. WGCNA: an R package for weighted correlation network analysis. BMC Bioinformatics. 2008;9: 559. doi:10.1186/1471-2105-9-559

15. Kuijjer ML, Tung MG, Yuan G, Quackenbush J, Glass K. Estimating Sample-Specific Regulatory Networks. iScience. 2019;14: 226–240. doi:10.1016/j.isci.2019.03.021

16. Ward JH. Hierarchical Grouping to Optimize an Objective Function. Journal of the American Statistical Association. 1963;58: 236–244. doi:10.1080/01621459.1963.10500845

17. Langfelder P, Zhang B, Horvath S. Defining clusters from a hierarchical cluster tree: the Dynamic Tree Cut package for R. Bioinformatics. 2008;24: 719–720. doi:10.1093/bioinformatics/btm563

18. White JD, Indencleef K, Naqvi S, Eller RJ, Hoskens H, Roosenboom J, et al. Insights into the genetic architecture of the human face. Nat Genet. 2021;53: 45–53. doi:10.1038/s41588-020-00741-7

19. Murtagh F, Legendre P. Ward’s Hierarchical Agglomerative Clustering Method: Which Algorithms Implement Ward’s Criterion? J Classif. 2014;31: 274–295. doi:10.1007/s00357-014-9161-z

20. Walakira A, Ocira J, Duroux D, Fouladi R, Moškon M, Rozman D, et al. Detecting gene– gene interactions from GWAS using diffusion kernel principal components. BMC Bioinformatics. 2022;23: 57. doi:10.1186/s12859-022-04580-7

21. WHO Global InfoBase team. The SuRF Report 2. Surveillance of chronic disease Risk Factors: Country-level data and comparable estimates. Geneva: World Health Organization; 2005.

22. Gillespie M, Jassal B, Stephan R, Milacic M, Rothfels K, Senff-Ribeiro A, et al. The reactome pathway knowledgebase 2022. Nucleic Acids Research. 2022;50: D687–D692. doi:10.1093/nar/gkab1028

23. O’Bray L, Rieck B, Borgwardt K. Filtration Curves for Graph Representation. Proceedings of the 27th ACM SIGKDD Conference on Knowledge Discovery & Data Mining. Virtual Event Singapore: ACM; 2021. pp. 1267–1275. doi:10.1145/3447548.3467442

24. Kruskal WH, Wallis WA. Use of Ranks in One-Criterion Variance Analysis. Journal of the American Statistical Association. 1952;47: 583–621. doi:10.1080/01621459.1952.10483441

25. Rand WM. Objective Criteria for the Evaluation of Clustering Methods. Journal of the American Statistical Association. 1971;66: 846–850. doi:10.1080/01621459.1971.10482356

26. R Core Team. R: A Language and Environment for Statistical Computing. Vienna, Austria: R Foundation for Statistical Computing; 2022. Available: https://www.R-project.org/

27. Fawcett KA, Barroso I. The genetics of obesity: FTO leads the way. Trends in Genetics. 2010;26: 266–274. doi:10.1016/j.tig.2010.02.006

28. The Prostate, Lung, Colorectal, and Ovarian (PLCO) Cancer Screening Trial, KORA, Nurses’ Health Study, Diabetes Genetics Initiative, The SardiNIA Study, The Wellcome Trust Case Control Consortium, et al. Common variants near MC4R are associated with fat mass, weight and risk of obesity. Nat Genet. 2008;40: 768–775. doi:10.1038/ng.140

29. D’Silva S, Chakraborty S, Kahali B. Concurrent outcomes from multiple approaches of epistasis analysis for human body mass index associated loci provide insights into obesity biology. Sci Rep. 2022;12: 7306. doi:10.1038/s41598-022-11270-0

30. Piñero J, Ramírez-Anguita JM, Saüch-Pitarch J, Ronzano F, Centeno E, Sanz F, et al. The DisGeNET knowledge platform for disease genomics: 2019 update. Nucleic Acids Research. 2019; gkz1021. doi:10.1093/nar/gkz1021

31. Zhu Z, Guo Y, Shi H, Liu C-L, Panganiban RA, Chung W, et al. Shared genetic and experimental links between obesity-related traits and asthma subtypes in UK Biobank. Journal of Allergy and Clinical Immunology. 2020;145: 537–549. doi:10.1016/j.jaci.2019.09.035

32. Storojeva I, Boulay J-L, Ballabeni P, Buess M, Terracciano L, Laffer U, et al. Prognostic and Predictive Relevance of DNAM-1, SOCS6 and CADH-7 Genes on Chromosome 18q in Colorectal Cancer. Oncology. 2005;68: 246–255. doi:10.1159/000086781

33. Kichaev G, Bhatia G, Loh P-R, Gazal S, Burch K, Freund MK, et al. Leveraging Polygenic Functional Enrichment to Improve GWAS Power. The American Journal of Human Genetics. 2019;104: 65–75. doi:10.1016/j.ajhg.2018.11.008

34. Lv H, Zhang M, Shang Z, Li J, Zhang S, Lian D, et al. Genome-wide haplotype association study identify the FGFR2 gene as a risk gene for Acute Myeloid Leukemia. Oncotarget. 2017;8: 7891–7899. doi:10.18632/oncotarget.13631

35. Spracklen CN, Karaderi T, Yaghootkar H, Schurmann C, Fine RS, Kutalik Z, et al. Exome-Derived Adiponectin-Associated Variants Implicate Obesity and Lipid Biology. The American Journal of Human Genetics. 2019;105: 15–28. doi:10.1016/j.ajhg.2019.05.002

36. Song M, Greenbaum J, Luttrell J, Zhou W, Wu C, Shen H, et al. A Review of Integrative Imputation for Multi-Omics Datasets. Front Genet. 2020;11: 570255. doi:10.3389/fgene.2020.570255

37. Lu Z. Canonical Correlation Analysis with Missing Values: A Structural Equation Modeling Approach. In: Wiberg M, Culpepper S, Janssen R, González J, Molenaar D, editors. Quantitative Psychology. Cham: Springer International Publishing; 2019. pp. 243–254. doi:10.1007/978-3-030-01310-3_22

38. Witten DM, Tibshirani RJ. Extensions of Sparse Canonical Correlation Analysis with Applications to Genomic Data. Statistical Applications in Genetics and Molecular Biology. 2009;8: 1–27. doi:10.2202/1544-6115.1470

39. Kriege NM, Johansson FD, Morris C. A survey on graph kernels. Appl Netw Sci. 2020;5: 6. doi:10.1007/s41109-019-0195-3

40. Zhang Y, Li T. Consensus clustering+ meta clustering= multiple consensus clustering. 2011.

41. Duroux D, Van Steen K. netANOVA: novel graph clustering technique with significance assessment via hierarchical ANOVA. bioRxiv. 2022 [cited 14 Feb 2023]. doi:10.1101/2022.06.28.497741

